# A Highly-ordered and Fail-safe Electrical Network in Cable Bacteria

**DOI:** 10.1101/866319

**Authors:** Raghavendran Thiruvallur Eachambadi, Robin Bonné, Rob Cornelissen, Silvia Hidalgo-Martinez, Jaco Vangronsveld, Filip J. R. Meysman, Roland Valcke, Bart Cleuren, Jean V. Manca

## Abstract

Cable Bacteria are an emerging class of electroactive organisms that sustain unprecedented long-range electron transport across centimeter-scale distances. The pathways of the electrical currents in these filamentous microorganisms remain unresolved. Here, the electrical circuitry in a single cable bacterium is visualized with nanoscopic resolution using Conductive Atomic Force Microscopy. Combined with perturbation experiments, it is demonstrated that electrical currents are conveyed through a parallel network of conductive fibers embedded in the cell envelope which are electrically interconnected between adjacent cells. This redundant structural organization of electrical pathways likely forms a crucial adaptive trait for organisms capable of long-distance electron transport. The observed electrical circuit architecture is unique in biology and could inspire future technological applications in bioelectronics.

---

Cable bacteria are filamentous microorganisms consisting of more than 10^4^ cells, forming unbranched filaments of up to several centimeters long.^[1,2]^ These bacteria thrive in both freshwater^[3,4]^ and marine sediments^[5–7]^, and couple the oxidation of sulfide within deeper strata of the sediment to the reduction of oxygen near the sediment-water interface. The spatial segregation of these redox half-reactions across centimeter distances necessitates electron transport, which occurs internally along the length of the filament.^[8–10]^ While nanometer-scale electron transport is known to occur in chloroplasts and mitochondria^[11–14]^, and micrometer-scale electrical currents are induced in the nanowire appendages of metal-reducing bacteria^[15–17]^, the centimeter-scale electron transport by cable bacteria extends the known length scale of biological transport by several orders of magnitude.^[2]^

While some aspects of the physiology and overall metabolism of cable bacteria have been recently resolved ^[18,19]^, the mechanism of long-distance electron transport (LDET) inside cable bacteria remains elusive. Scanning Electron Microscopy (SEM) and Atomic Force Microscopy (AFM) have shown that the surface of cable bacteria has a unique topography, with parallel ridges running along the entire length of the filaments.^[8,20,21]^ At the junction between two cells, a ‘cartwheel’ structure (**Figure 1A**) is observed^[20]^, of which the composition and function is still unknown. Initial Electrostatic Force Microscopy (EFM) experiments have shown a distinct elevation of electrostatic forces over these ridges, suggesting the presence of underlying polarizable structures that possess a charge storage capacity.^[8]^ Subsequent electron microscopy imaging has revealed a parallel network of fibers embedded within the periplasm underneath the ridges. This periplasmic fiber network can be selectively extracted from the cell envelope of the cable bacteria^[20]^, thus retaining a so-called fiber sheath, and electrical probe measurements have demonstrated that this fiber sheath harbors conductive material enabling long-range electron transport in cable bacteria.^[22]^ The parallel fibers make up the majority of the biomass material in the fiber sheath, and so the hypothesis has been coined that the periplasmic fibers are the conductive pathways within cable bacteria.^[22]^

**Figure 1.**
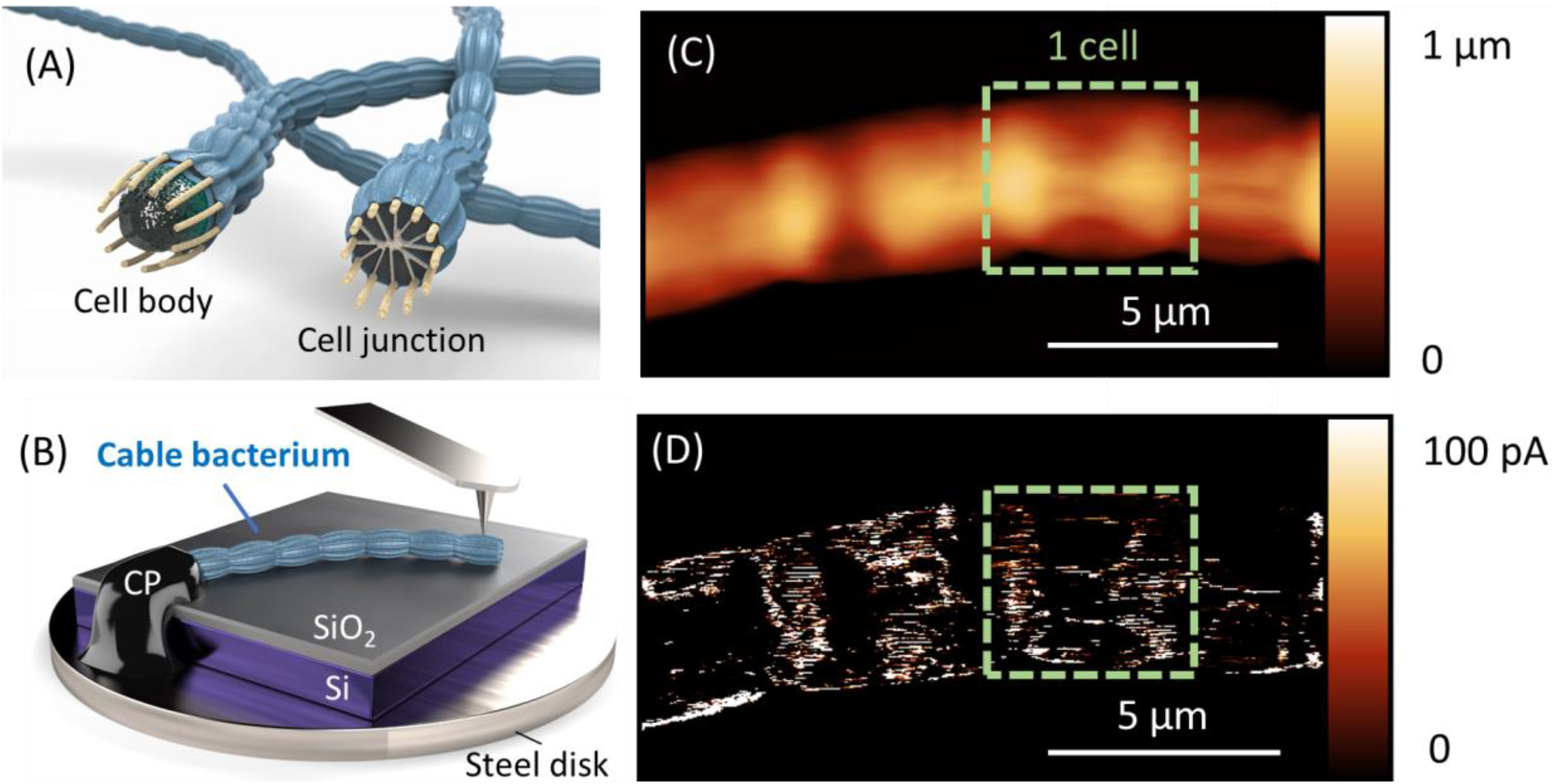
C-AFM measurements visualize parallel current pathways in a cable bacterium (A) Geometric model: a cable bacterium is a filament of thousands of cells containing fibers that run continuously in the periplasmic space along the filament. ^[20]^ In the junction between two cells, a ‘cartwheel’ structure is present, for which the function and composition is unknown. (B) A single cable bacterium filament is placed on a silicon (Si) substrate with silicon oxide (SiO_2_) top layer for conductive AFM imaging. The AFM functions as a two electrode system, with one end of the bacterium connected via carbon paste (CP) to a steel disk, while the other end is probed with the AFM tip. Simultaneously with obtaining (C) a height profile, a potential is applied between the two electrodes, for which (D) the current response is measured. Parallel lines show that currents are flowing through periplasmic fibers upon direct contact of the AFM tip.

In order to precisely pinpoint the location and connection of the conductive pathways within cable bacteria, we employed Conductive Atomic Force Microscopy (C-AFM) at nanoscale resolution. This technique enables to measure simultaneously the topography of a material together with the electric current flow at the contact point with the surface, and has been previously used to demonstrate conductance of the surface appendages of metal-reducing bacteria^[15,23]^, and cyanobacteria.^[24]^ In addition, we designed dedicated perturbation experiments, where we deliberately interrupted the conductive pathways in order to investigate the electrical robustness of damaged cable bacteria with respect to long-distance electron transport.

In a first experiment, an intact filament was isolated from a sediment enrichment culture and placed on a non-conductive SiO_2_ substrate in the C-AFM setup under N_2_ atmosphere (see materials and methods). At one end of the filament, an electrical connection was established with conductive carbon paste, whereas the other end was left free for imaging with the C-AFM probe (**Figure 1B**). The C-AFM setup thereby functions as a two-electrode system: when a voltage is applied between the (movable) probe and the (fixed) electrical contact, the detection of a current indicates the presence of a conductive pathway within the cable bacterium filament. Initial C-AFM measurements on intact filaments were performed using soft SCM-PIT-V2 probes (see experimental section). A first scan of a certain area revealed almost no conductive structure, indicating that the conductive pathways cannot be directly contacted at the surface of the cable bacterium (Figure S1A,B). After consecutive scanning of the same area, only a few new tiny areas of current pathways appeared (Figure S1C,D), suggesting that conductive structures were embedded underneath the surface, covered by a thin insulating layer.

When we adapted our scanning procedure and used a stiff diamond coated CDT-NCLR probe (see experimental section), a clear electrical response pattern became apparent from the very first scan (**Figure 1C,D**). Parallel conducting lines were observed along the length of the filament, spatially separated from each other by non-conductive zones. After only two scans, the current at a given area vanished again. This suggests that the stiff AFM tip opens up the insulating outer layer at the first scan, but keeps on removing material in subsequent scans, until also the conductive fibers are etched away. This gradual removal of material was confirmed in consecutive images: after about 10 consecutive scans the entire cable bacterium filament was sheared off and abraded material was deposited outside the imaged area (Figure S2). A course calibration of the AFM height data (10 abrasive scans remove ~ 500 nm height of cable bacterium material) indicates that the conductive fiber structures must be contained within a <100 nm narrow zone, which agrees well with previous fiber thickness characterizations (~50 nm).^[20]^

To demonstrate that the periodicity of the observed conductive pattern coincides with the previously reported fiber network in the cell envelope of the cable bacteria, we subjected filaments to an SDS/EDTA extraction procedure that removes the cytoplasm and membranes.^[20]^ This extraction leaves behind a so-called fiber sheath, which embeds the regularly spaced periplasmic fibers (diameter ~ 50 nm, with an interspacing of ~ 200 nm) in a rigid basal sheath. Recently, macroscopic electrical measurements have shown that this fiber sheath endows the conductivity upon cable bacterium filaments, but no definite proof was given that the fibers are truly guiding the electrical currents.^[22]^

When fibers sheaths were subjected to the C-AFM scanning procedure with the stiff diamond probe, the parallel pattern of conductive pathways was again observable (Figure S3). In order to correlate electrical current with height measurements, we performed scans with the soft SCM-PIT-V2 probe for a new sample (**Figure 2A,B**). In a transect perpendicular to the fiber direction, conductive zones (133 ± 24 nm wide) were separated by non-conductive interspacing (62 ± 8 nm wide) (**Figure 2C**). The zones where a current was measured corresponded to the elevated regions caused by the fibers (Figure 2C), with a peak-to-peak width for the current profile (194 ± 20 nm) corresponding to that of the height profile (191 ± 19 nm) and the fiber interspacing known from literature.^[20]^ These results unequivocally demonstrate that the periplasmic fibers are the conductive structures in cable bacteria.

**Figure 2.**
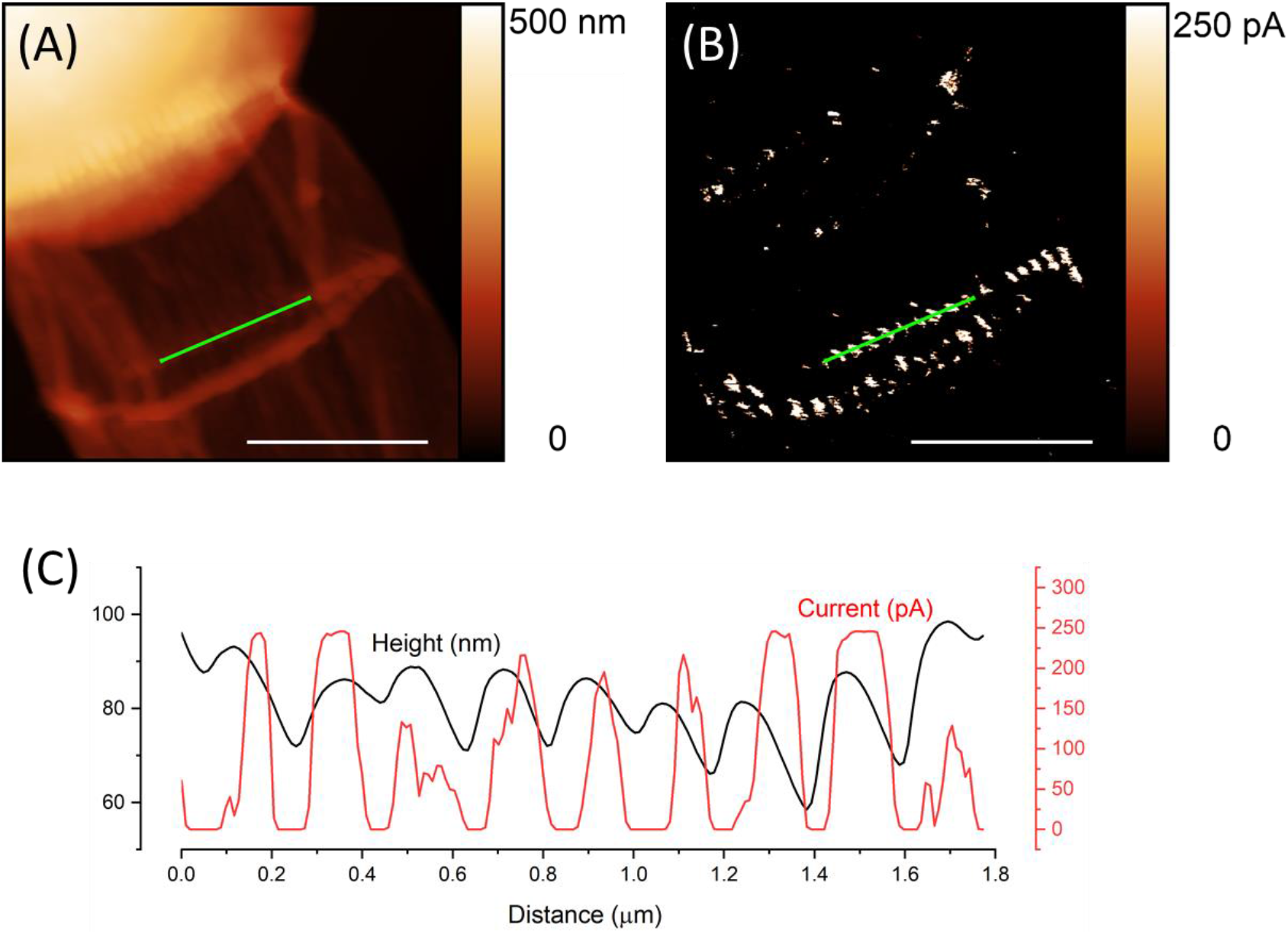
Correlation of ridges with conductive structures. (A) Height and (B) current profiles were obtained for a fiber sheath with a soft AFM probe. (C) A transect line profile for the green line in the images (A) and (B). The location of the fibers (maxima in the height profile; black curve) corresponds to locations with zones of increased current flow (red curve).

Cable bacteria hence contain a set of parallel conductive fibers that can transport electrons over centimeter distances along the longitudinal axis of the filament. However, if fibers would act as independent conductive pathways, the resulting electrical network would be highly vulnerable to disruption. Any incidental damage to a fiber would immediately annihilate the conductive fiber channel along the entire length of the filament. Hence, one would expect an evolutionary drive towards an electrical connectivity between fibers. Detailed electron microscopy imaging has previously shown that the junction between two cells in a filament contains a ‘cartwheel’ structure (Figure 1A), which starts from each fiber and radiates inwards to a central node.^[20]^ The function of this cartwheel structure is currently unknown, but it might function as an electrical interconnection between the fibers, thus equipping the electrical grid with a redundant and safety-critical topology.

In order to evaluate whether fibers are electrically connected at the cell junctions, a perturbation experiment was designed in which we deliberately “cut” the network in specific locations via abrasive removal of material within a narrow zone during sequential contact mode imaging with C-AFM (**Figure 3**). When such cuts were made in two consecutive cells of a filament, the whole area surrounding the cuts retained a conductive pathway to the downstream electrode (Figure 3A-C). However, when the cuts were made within the same cell, so that all parallel fibers were cut at least once, only the area downstream from the two cuts showed a conductive connection to the electrode (Figure 3D-F). These results demonstrate that the fibers in the cell envelope are not electrically connected within the cell body, but the cell junctions do contain an electrical interconnection, which is most likely due to the conductive cartwheel structure (Figure 1A).

**Figure 3.**
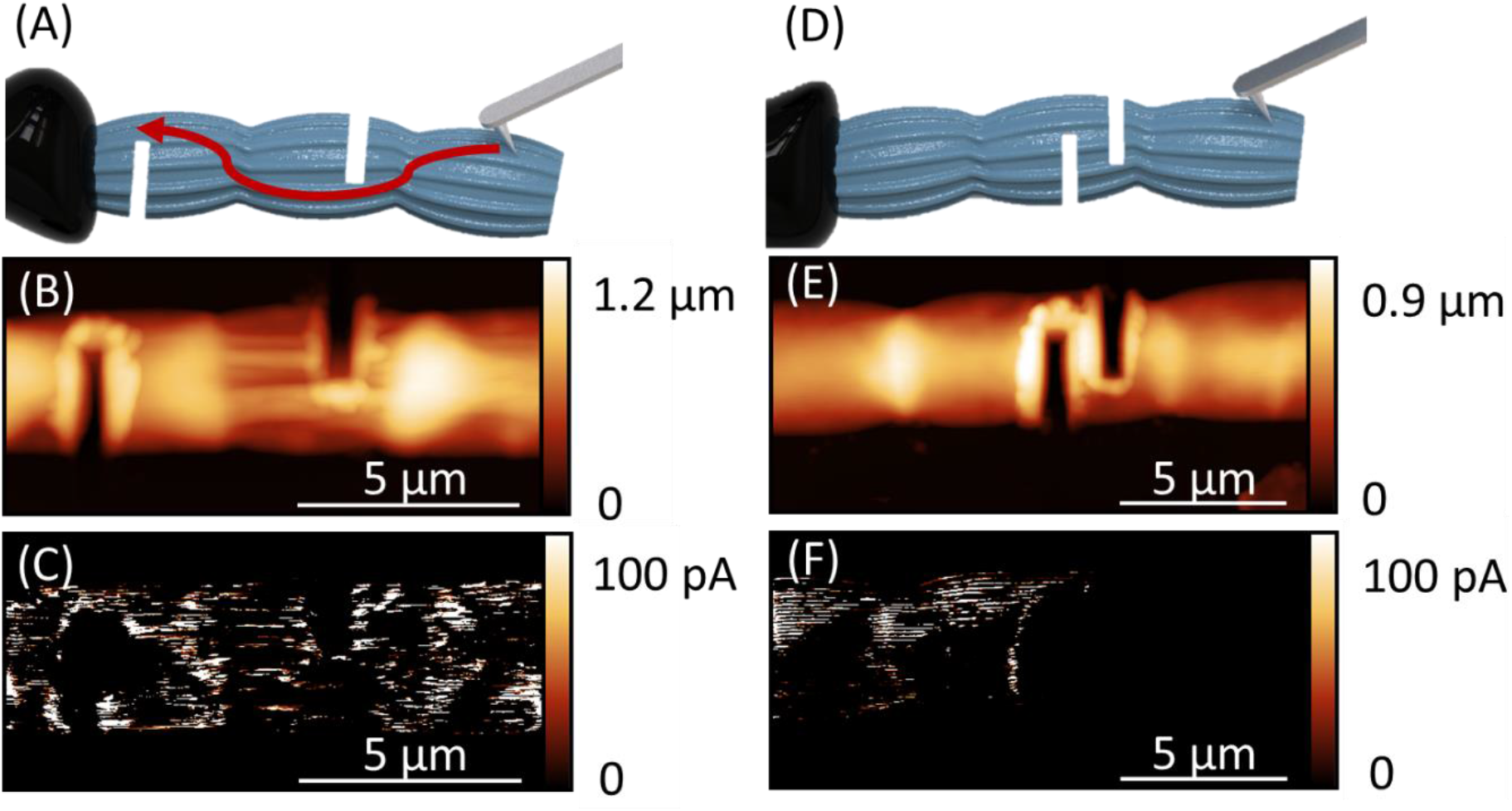
C-AFM measurements show an electrical interconnection of the fibers at cell junctions. (A) Two cuts are made in two adjacent cells of a filament after which (B) the height and (C) the current are imaged with C-AFM. Current flowing from the left side through all fibers to the carbon paste (CP) electrode proves an interconnection of the fibers. When a cut is made in a single cell (D-E), the current vanished completely on the right side (F), thereby proving the interconnection to be present in the cell junctions.

In this work we show that cable bacteria have evolved a conductive network with a scale and a degree of structural organization that is unprecedented in nature. Our C-AFM measurements reveal that centimeter-scale electrical currents are transported through a parallel network of conductive periplasmic fibers, which is redundantly interconnected at the cell junctions. Such an interconnected wiring configuration generates electrical robustness, which most likely provides an evolutionary advantage, as it maintains its electron transport system in the presence of local defects or cuts. This demonstrates that nature has already invented highly ordered and fail-safe electrical networks long before the era of microelectronics.

## Experimental Section

### Cultivation and specimen preparation

Sediments containing cable bacterium filaments were collected from the creek bed of a salt marsh (51°26’21.6“N 4°10’08.3”E), which has previously been reported to grow cable bacteria.^[5]^ Cable bacterium filaments were enriched in laboratory sediment cores under oxygenated overlying water, isolated from the sediment and washed to remove sediment particles and attached bacteria, using the procedure as detailed in ^[20]^. This produces so-called “intact” filaments. Fibers sheaths were obtained from intact filaments using the sequential extraction procedure as described in ^[20]^ and ^[22]^.

For the C-AFM measurements, 1cm × 1cm diced Si wafers with a SiO_2_ top layer (200 nm thick) were mounted on stainless steel discs (12 mm diameter) using silver paste as an adhesive. A single cable bacterium filament was placed on the substrate with glass hooks, either as an intact filament or a fiber sheath. An aqueous graphene dispersion (henceforth referred to as “carbon paste”, EM-Tec C30 Micro to Nano, Haarlem, the Netherlands) was applied over one end of the cable bacterium filament. One of the ends was electrically connected to the steel disc using carbon paste (Figure 1). The samples were stored for two minutes in the load lock of the glove box under vacuum for the carbon paste to dry, and finally placed under the AFM.

### Conductive Atomic Force Microscopy

C-AFM was performed using Bruker (Santa Clara, CA, USA) Multimode 8 AFM with a Nanoscope V controller. The AFM was housed within a home-made glove box system under a nitrogen atmosphere to prevent the decay of conductance of the bacterial filaments resulting from exposure to O_2_.^[22]^ Two types of electrical probes were used. In one set of measurements, we used Conductive Diamond coated Tip – Non-Contact/tapping mode – Long cantilever – Reflex coating (CDT-NCLR) probes supplied by NanoWorld AG (Neuchâtel, Switzerland), containing a highly doped diamond coated tip (spring constant 70 N/m). These probes have a macroscopic tip radius between 100 nm and 200 nm and are extremely wear resistant due to the diamond coating. Alternatively, we also used SCM-PIT-V2 probes (Bruker, spring constant 3 N/m), where the front side of the probe was coated with Pt-Ir (apex radius 25 nm).

The probe holder was electrically connected to a Peakforce Tunnelling AFM (PF-TUNA) module, which in turn was connected to the Nanoscope V controller. Sample voltage bias was applied through the magnetic sample holder, and a standard sample bias of 0.5V was applied. The height images and current images have the same spatial resolution, since both data types are measured for every pixel. The microscope was used in Scanasyst mode while localizing an area of interest to prevent unintentional damages and to obtain high resolution topography images, before switching to conductive contact mode. Finally, AFM data were analysed using the open source SPM analysis software Gwyddion.

## Acknowledgements

We thank our colleagues from X-LAB from Hasselt University and the Microbial Electricity team from University of Antwerp for discussions and feedback. Special thanks to H. T. S. Boschker, I. Cardinaletti, J. Drijkoningen, JL. Hou and S. Thijs for their insights and discussions. Thanks to K. Ceyssens and T. Custers for the graphics.

This research was financially supported by the Research Foundation - Flanders (FWO project grant G031416N to FJRM and JM and FWO aspirant grant 1180517N to RB).

All measurements and data analysis were performed by RTE and RB in equal contribution. JM and BC coordinated the study. Conceptualization and discussion were done by JM, RV, BC, RC, RTE and RB. Cable bacteria enrichment and fiber sheath extraction was performed by SHM, RB and FJRM. Funding was acquired by JM, JV and FJRM. Writing was done by RB, RTE, JM, BC and FJRM, with contributions from all authors.

