## Supplementary information for "A Highly-ordered and Fail-safe Electrical Network in Cable Bacteria"

Stability in nitrogen atmosphere

To check the possible decay of intact filaments in nitrogen atmosphere, we measured the resistance of single filaments using the ramp mode immediately after introducing the sample to the AFM. Doing the same ramp measurement more than 65 hours later, a slight increase in resistance of only 3.4% was recorded.

C-AFM analysis of intact filaments using a soft probe

An area for an intact filament was imaged using the soft SCM-PIT-V2 probe as shown in Figure S1. Figure S1A is the first height image of the of this area, with Figure S1B showing not much current for this area. After 27 consecutive scans of the same area, Fig S1C shows no significant change in topography. However, a few other but insignificant areas of current appeared (Fig S1D).

Removal of material

We show that with the use of CDT-NCLR probes there is a rapid material removal due to the abrasive nature of the coated diamond tips. For one experiment, we measured an area on a continuous basis. Figure S2A is the Scanasyst topography image of an area after a cut was made in the filament, with Figure S2B being the first contact mode image. Figure S2C is the fifth in series, just before the filament was broken into two, and finally, Figure S2D is the remnant of the filament (13th image). Removed material was located just outside of the scan area Figure S2E.

C-AFM analysis of fiber sheaths using a hard probe

Previous work^[17]^ has shown that the fiber sheath contains the conductive structures of the cable bacterium filament. Since the detergent treatment removes the outer membrane, the conductive fibers is expected be easy to reach with electrically conductive AFM probes. Fiber sheaths imaged using CDT-NCLR probes reveal the conductive pathways within the ridge structures (Figure S3). A current profile is found from the very first scan.

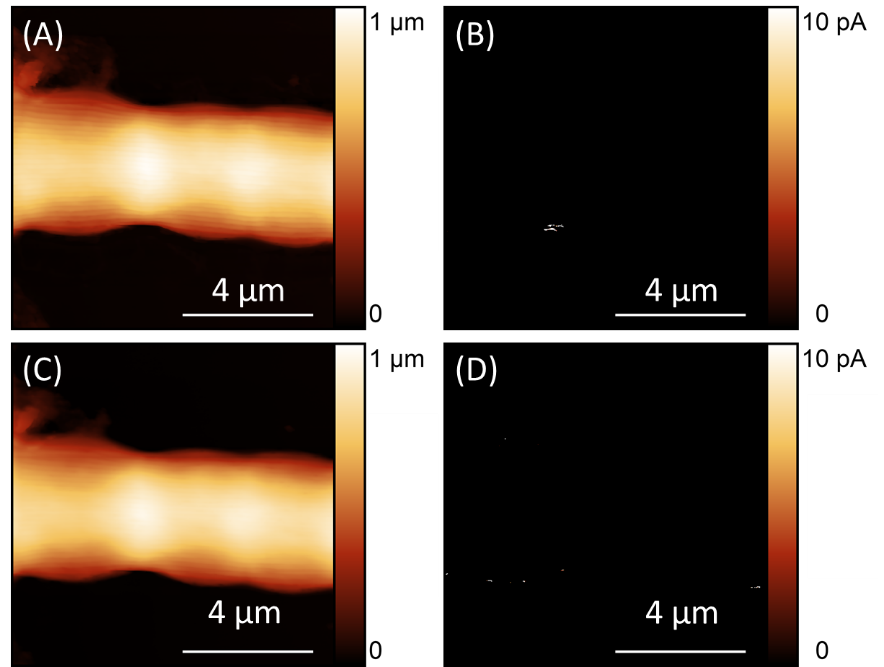

**Figure S1.** C-AFM at the junction of an intact filament using SCM-PIT V2 probe. Image (A) was taken at the start of the measurement series, for which (B) the current is shown. The imaging was set to continuous capture. (C) After 27 scans were performed, the current (D) showed only a few new but insignificant areas to be conductive, but no clear current profile could be drawn from it.

5 µm

5 µm

5 µm

5 µm

5 µm

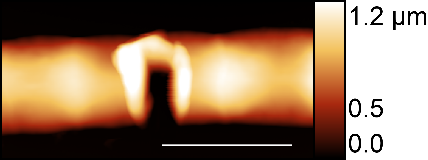

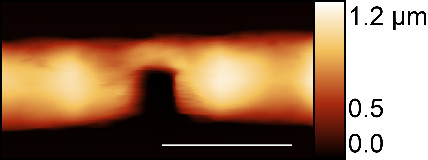

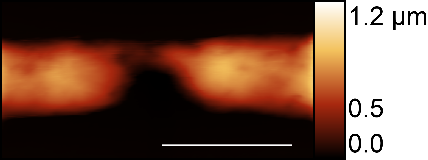

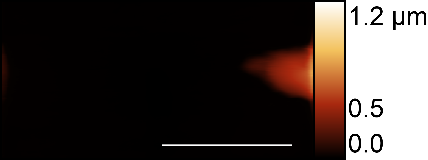

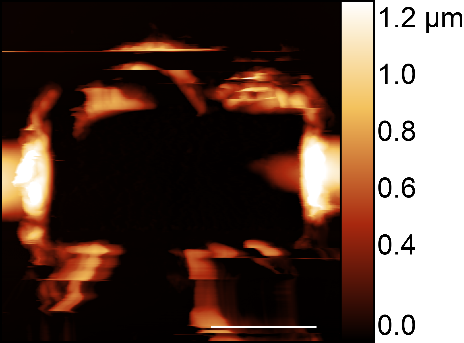

(A)

(B)

(C)

(D)

(E)

**Figure S2.** Gradual removal of material using a hard CDT-NCLR probe. (A) Height image using Scanasyst mode after creating a cut. Contact mode C-AFM images (B) first, (C) 5th and (D) 13th. (E) is the Scanasyst image of the enlarged area showing the debris collected just outside the scan area.

5 µm

5 µm

(B)

(A)

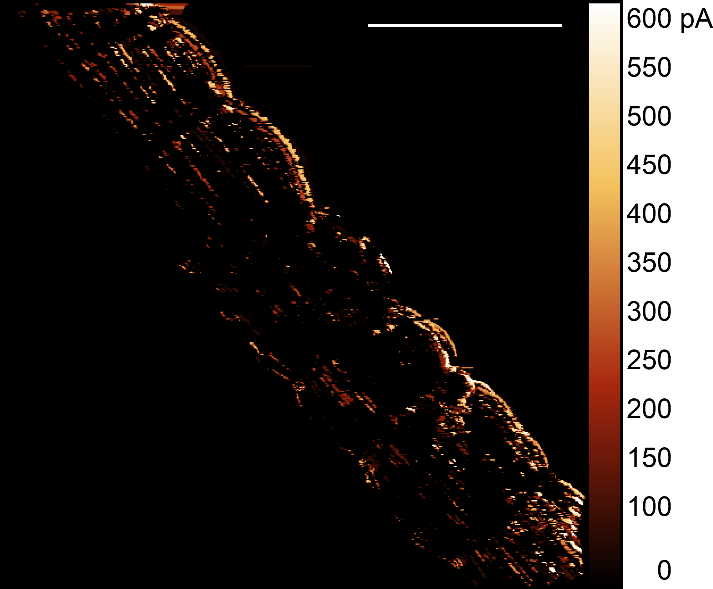

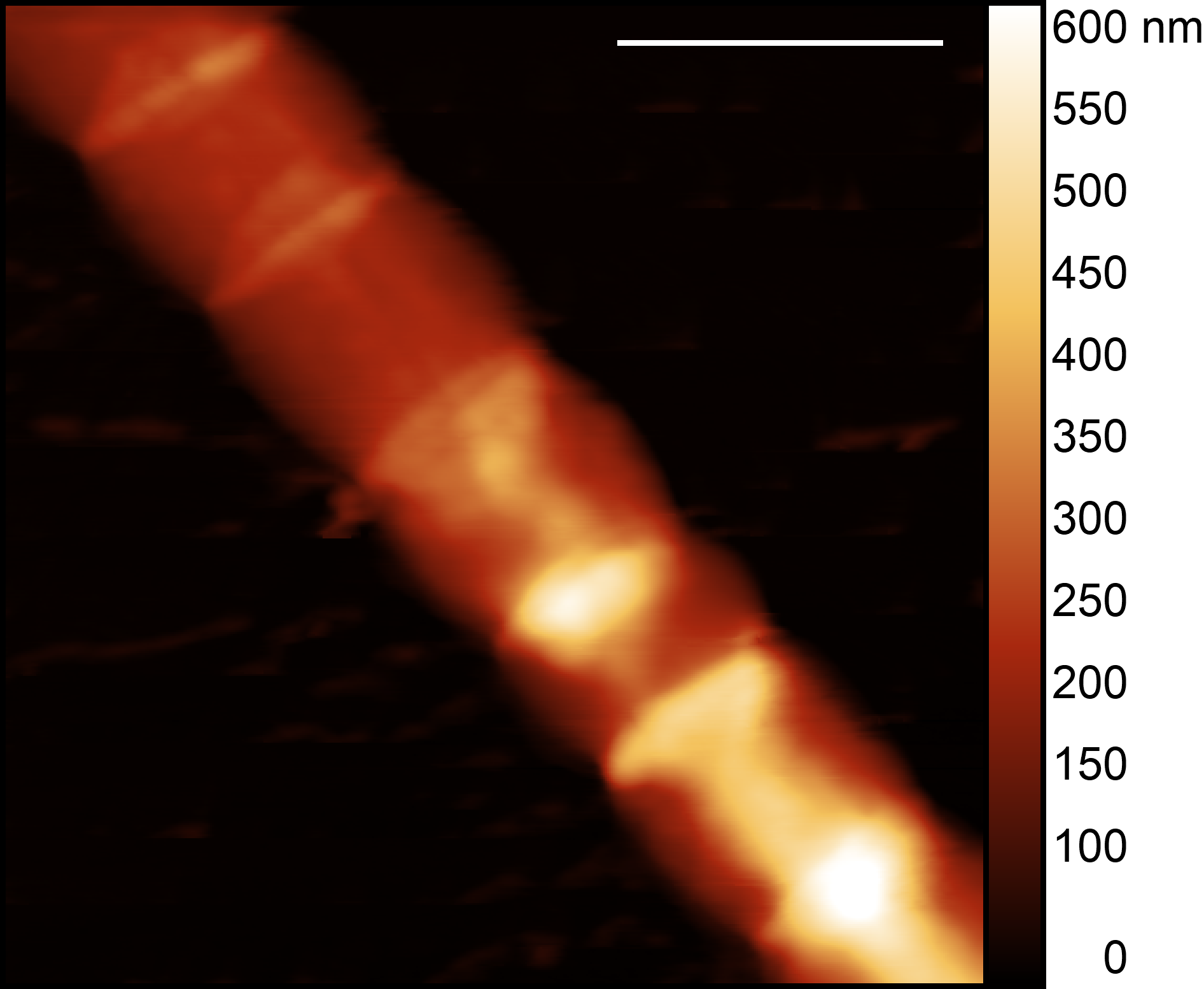

**Figure S3.** C-AFM measurements of a fiber sheath (A) Height and (B) current images of a fiber sheath imaged using CDT-NCLR probe.
